# Self-immunity guided identification of threonyl-tRNA synthetase as the molecular target of obafluorin, a *β*-lactone antibiotic

**DOI:** 10.1101/704981

**Authors:** Thomas A. Scott, Sibyl F. Batey, Patrick Wiencek, Govind Chandra, Silke Alt, Christopher S. Franklyn, Barrie Wilkinson

**Author notes:** These authors contributed equally to this work (Thomas A. Scott, Sibyl F. Batey).

## Abstract

To meet the ever-growing demand of antibiotic discovery, new chemical matter and antibiotic targets are urgently needed. Many potent natural product antibiotics which were previously discarded can also provide lead molecules and drug targets. One such example is the structurally unique *β*-lactone obafluorin, produced by *Pseudomonas fluorescens* ATCC 39502. Obafluorin is active against both Grampositive and -negative pathogens, however the biological target was unknown. We now report that obafluorin targets threonyl-tRNA-synthetase and we identify a homologue, ObaO, which confers self-immunity to the obafluorin producer. Disruption of *obaO* in *P. fluorescens* ATCC 39502 results in obafluorin sensitivity, whereas expression in sensitive *E. coli* strains confers resistance. Enzyme assays demonstrate that *E. coli* threonyl-tRNA synthetase is fully inhibited by obafluorin, whereas ObaO is only partly susceptible, exhibiting a very unusual partial inhibition mechanism. Altogether, our data highlight the utility of a self-immunity guided approach for the identification of an antibiotic target *de novo* and will ultimately enable the generation of improved obafluorin variants.

## Introduction

*β*-Lactone rings occur infrequently in Nature but are constituents of several different natural product classes.^1^ Structurally similar to *β*-lactam rings, they are effective electrophiles able to form covalent linkages with nucleophilic residues of target proteins^2^ and possess significant therapeutic value as hydrolase inhibitors.^3^ Prominent examples include the hybrid polyketide-non-ribosomal peptide (PK-NRP) Salinosporamide A (marizomib), which is a potent 20S proteasome inhibitor that is currently in Phase III clinical trials as a treatment for glioblastoma,^4^ and tetrahydolipstatin (orlistat), a semi-synthetic derivative of the PK-NRP lipstatin that is approved by the US Food and Drug Administration for the treatment of obesity due to its powerful lipase inhibitory activity.^5^ Further validated targets of natural and synthetic *β*-lactone inhibitors include serine hydrolase and polyketide synthase enzymes associated with mycolic acid biosynthesis, inhibition of which is lethal for *Mycobacterium tuberculosis*,^6^ as well as esterases, cutinases, homoserine transacetylase, cathespin A, ClpP^7^ and *N*-acylethanolamine acid amidase.^8^

Due to their significant therapeutic potential, much effort has been put into elucidating the genetic and biochemical basis for *β*-lactone ring formation during natural product biosynthesis.^9^ Recent work in our lab^10^ and by others^11^ characterized the biosynthetic pathway to the *β*-lactone antibiotic obafluorin (**1**) from *Pseudomonas fluorescens* ATCC 39502. **1** is assembled by ObaI, a bimodular non-ribosomal peptide synthetase (NRPS) which catalyzes amide bond formation between 2,3-dihydroxybenzoic acid (**2**), and the nonproteinogenic amino acid (2*S*,3*R*)-2-amino-3-hydroxy-4-(4-nitrophenyl)butanoate (**3**), the product of a rare L-threonine transaldolase (Fig. 1). Critically, it is this unusual *β*-hydroxy-*α*-amino acid precursor that comprises the functional groups which give rise to the *β*-lactone ring of **1**, following cyclization catalyzed by a rare type I thioesterase domain present in ObaI, which possesses a noncanonical active site Cys residue.

**Fig. 1.**
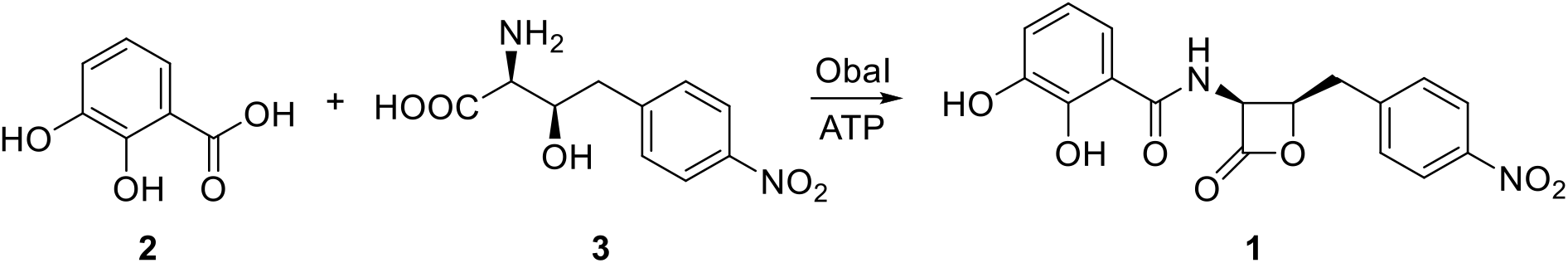
Biosynthesis of obafluorin (1) in *P. fluorescens* ATCC 39502.

Despite undergoing facile hydrolysis and ring opening in the presence of nucleophiles, **1** exhibits potent antibacterial activity against a range of Gram-positive bacteria and is also active against Gram-negative bacteria including *E. coli* and various *Pseudomonas aeruginosa* strains (Table 1 and ESI Fig. 1).^12,13^ When dosed systemically, it protects mice challenged with *Streptococcus pyogenes* (ED_50_ = 50 mg/kg) and is also reported to cause an unusual cell-elongation phenotype in *E. coli* cells exposed to sub-lethal doses. These various observations suggest that **1** acts in a specific manner rather than as a general acylating agent, yet, despite this unusual activity profile, the molecular target for **1** remained obscure.

Given the urgent need for new antibacterial targets for antibiotic development, we turned our attention to identifying the molecular target of **1**.

## Results and discussion

### Bioinformatic analysis identifies a putative 1 resistance gene

We began our investigation by examining the genomic neighborhood surrounding the *oba* biosynthetic gene cluster (BGC) (MiBIG accession: BGC0001437; GenBank accession: KX931446.2) in search of self-resistance gene candidates. Genes encoding immunity determinants commonly co-occur with the biosynthetic loci required for the production of natural product antibiotics,^14^ a phenomenon that has inspired recent genome-mining efforts to identify novel scaffolds with known targets.^15–17^ Comparative genomic analysis of the *oba* BGC characterized in *P. fluorescens* ATCC 39502 with other putative **1** BGCs present in the genomes of two *P. fluorescens* soil isolates, several *Burkholderia* species and *Chitiniphilus shinanonensis* DSM 23277 (ESI Table 1) allowed the identification of a minimal set of genes conserved across all clusters (Fig. 2). In addition to those previously ascribed biosynthetic or regulatory functions,^10^ only one additional gene, encoding a putative threonyl-tRNA synthetase (ThrRS), was present in all putative **1** BGCs; we have named this gene *obaO* (GenBank accession: KX931446.2). Aminoacyl-tRNA synthetases (aaRSs) play an essential role in protein synthesis, specifically activating and loading amino acids onto their cognate tRNAs.^18^ Although ubiquitous, divergence in aaRS sequence structure has led to the evolution of NP antibiotics that selectively target those of competing organisms.^19,20^ In a number of instances, a self-resistance isoform of the target aaRS is encoded within the NP BGC as an addition to the primary housekeeping copy encoded elsewhere in the genome of the producing-organism.^21–23^ Indeed, the putative *oba* BGC-associated ThrRS homologues all represent second copies in their respective genomes and, likewise, an additional copy of ThrRS is only observed in the presence of the *oba* BGC amongst these genera.

**Fig. 2.**
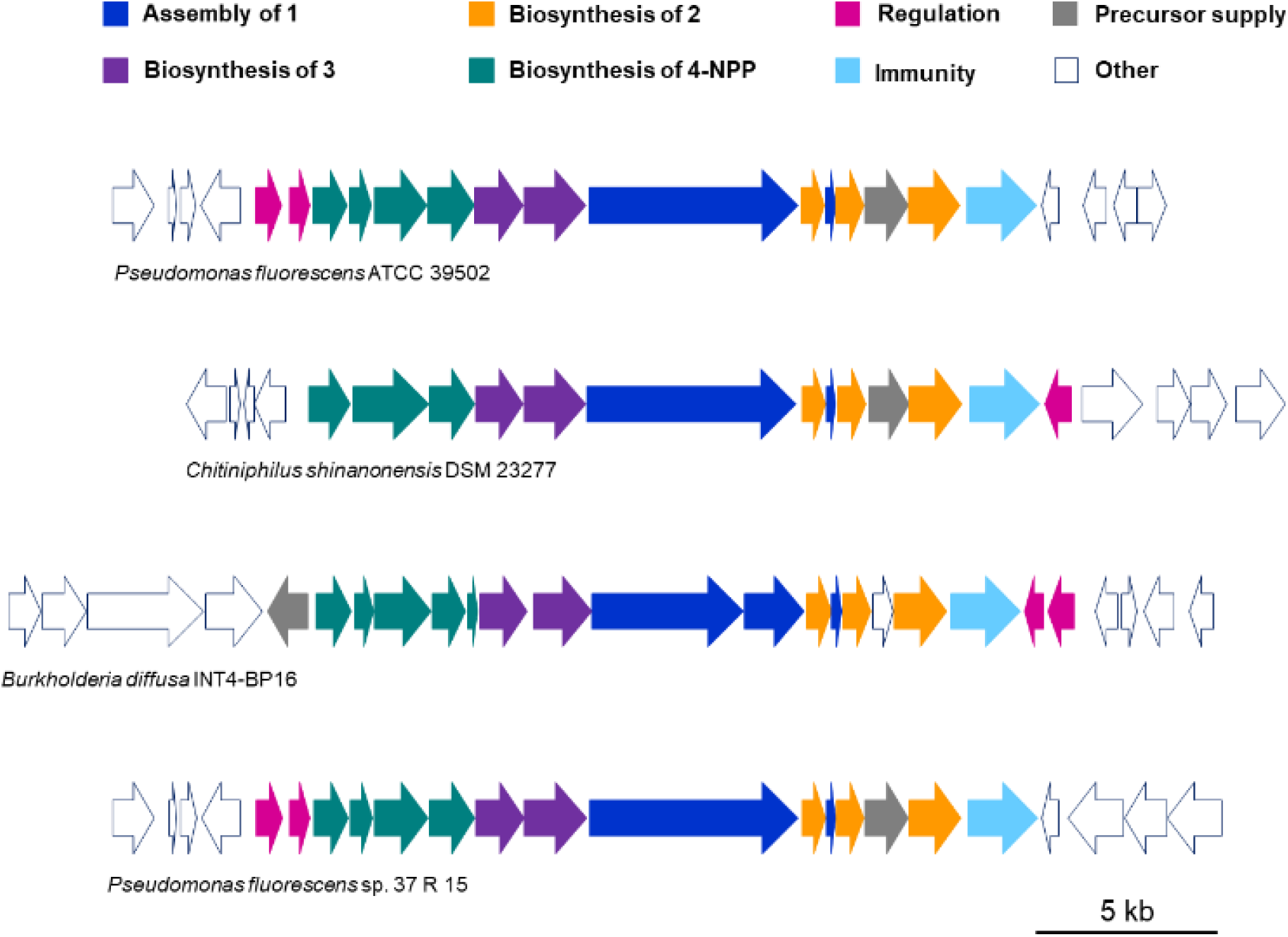
Comparative analysis of 1 biosynthetic gene clusters. Genes are colour-coded according to their function, with putative resistance genes highlighted in cyan. 4-NPP = 4-Nitrophenylpyruvate, an intermediate in the biosynthesis of **3**.

### *obaO* confers self-immunity to 1 in the native producer

To confirm the ability of ObaO to confer **1** immunity in the native producer, we attempted *obaO* inactivation experiments using the pTS1 suicide vector generated previously to perform knockouts in biosynthetic *oba* genes.^10^ However, following double crossover and counter-selection, we observed that resulting colonies were always exclusively wild-type (WT), consistent with a selection pressure against losing the immunity determinant when the *oba* BGC is functional.

To circumvent this issue, we deleted *obaO* in a Δ*obaL* background in which **1** production is abolished but can be rescued by exogenous supplementation with **2**. In the absence of exogenous **2** the growing strains accumulate increased levels of the shunt metabolites 4-nitrophenylethanol and 4-nitrophenylacetate, which is indicative of an otherwise functional biosynthetic pathway.^10^ We were successful in generating the Δ*obaL*Δ*obaO* strain and resulting mutants were verified by PCR amplification across the newly deleted region, with subsequent sequencing of the amplicon. When the Δ*obaL*Δ*obaO* strain was grown in production medium supplemented with **2**, growth was strongly inhibited, in contrast to cultures of WT and Δ*obaL* supplemented with **2**, and un-supplemented controls (Fig. 3). The Δ*obaL*Δ*obaO* strain was subsequently complemented genetically by ectopic expression of *obaO* using the pJH10TS vector described previously,^10^ which restored WT-like growth and **1** production when the strain was grown in **2**-supplemented production medium (Fig. 3). Taken together, these observations are consistent with a role for ObaO in conferring self-immunity to **1** in the native producer, *P. fluorescens* ATCC 39502.

**Fig. 3.**
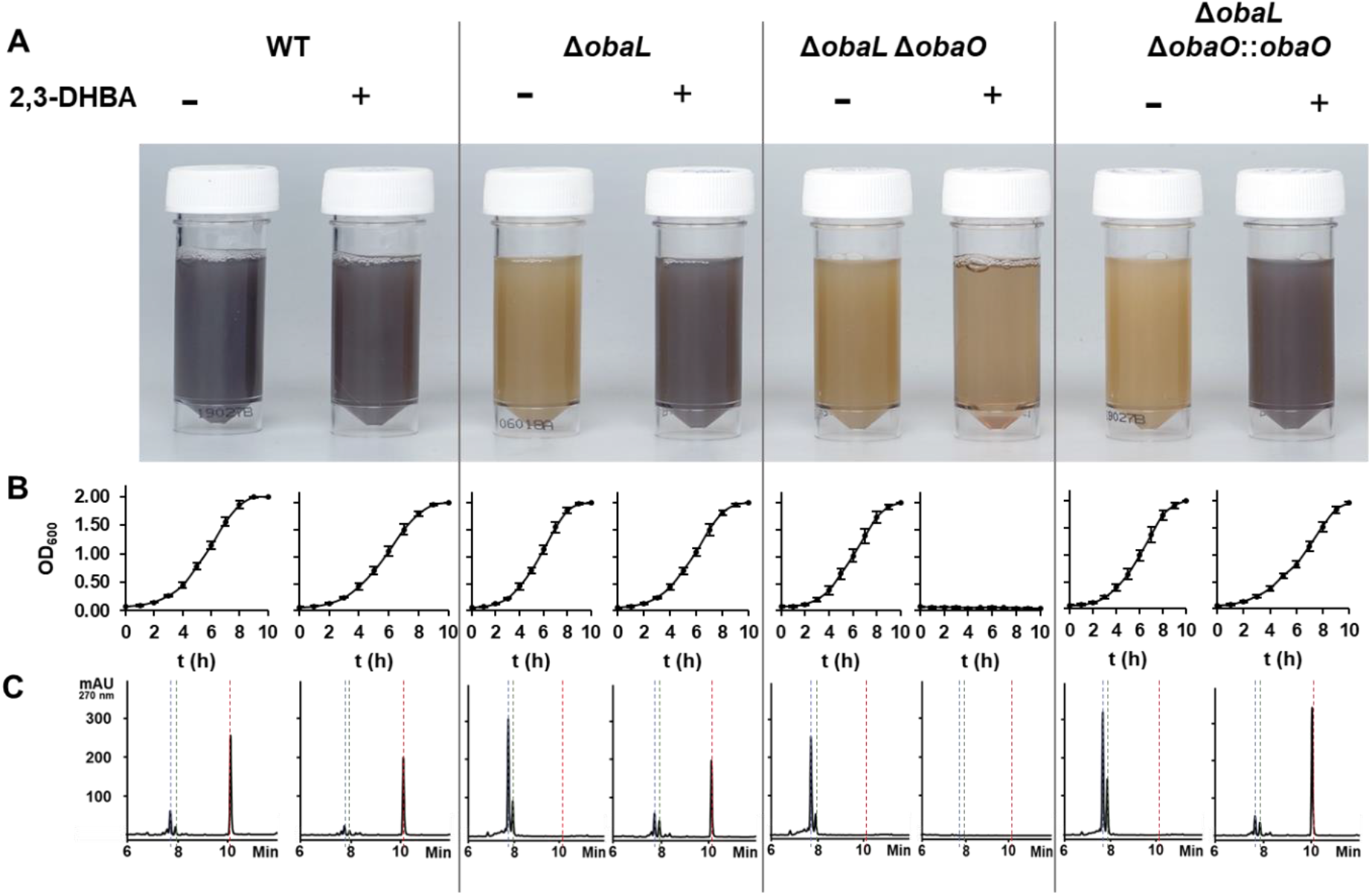
ObaO is the immunity determinant for 1 in *P. fluorescens* 39502. WT, Δ*obaL*, Δ*obaL*Δ*obaO* and Δ*obaL*Δ*obaO::obaO* strains were grown ± **2** (0.2 mM). As shown previously,^10^ **2** restores **1** production for Δ*obaL*, whereas the addition of **2** to Δ*obaL*Δ*obaO* abolishes growth. **A)** Aliquots of each strain after 14 h growth, with the purple colouration being indicative of **1** production. **B)** Log phase growth curves, showing complete absence of growth for Δ*obaL*Δ*obaO* + **2**. Each data point is the average of three biological repeats and bars show the standard error. **C)** Representative HPLC chromatograms at 270 nm for each condition (taken at 14 h for each of the three biological replicates); **1** elutes at 10.1 min (red dashed line) and the shunt metabolites 4-nitrophenylethanol (blue dashed line) and 4-nitrophenylacetate (green dashed line) elute at 7.7 min and 7.9 min respectively. These accumulate in growing strains where the biosynthesis of **2** is disrupted but the biosynthetic machinery is otherwise intact.^10^

### *obaO* confers transferable high-level immunity to 1-sensitive *E. coli* strains

To further validate our hypothesis, we introduced the *obaO* gene into **1**-sensitive bacterial strains to investigate its ability to confer resistance upon those strains when challenged with **1**. Given that *E. coli* was previously identified as being sensitive to **1**,^12^ we tested the clinical strain *E. coli* ATCC 25922 and *E. coli* NR698, which has increased permeability to antibiotics due to compromised outer-membrane lipopolysaccharide assembly.^24^ Off-target reactions of **1** with culture medium components precluded the use of microbroth assays, so we turned to disc-diffusion based approaches. We also found that **1** binds to paper discs, as reported previously,^25^ so we instead employed a spot-on-lawn approach by directly applying a solution of **1** dissolved in acetonitrile to the surface of agar bioassay plates. Using this method, **1** was shown to have a minimum inhibitory concentration (MIC) of 256 μg/mL for *E. coli* 25922, and 4 μg/mL for *E. coli* NR698 (Fig. 4, Table 1 and ESI Fig. 1). However, when *obaO* was expressed ectopically in both strains using the pJH10TS vector, both became resistant to **1** up to the highest concentration tested of 2000 μg/mL. This represents increases in the MIC of **1** of at least 8- and 250-fold for *E. coli* ATCC 25922 and *E. coli* NR698, respectively, relative to empty vector controls. Therefore, *obaO* confers transferable high-level immunity to **1**.

**Fig. 4.**
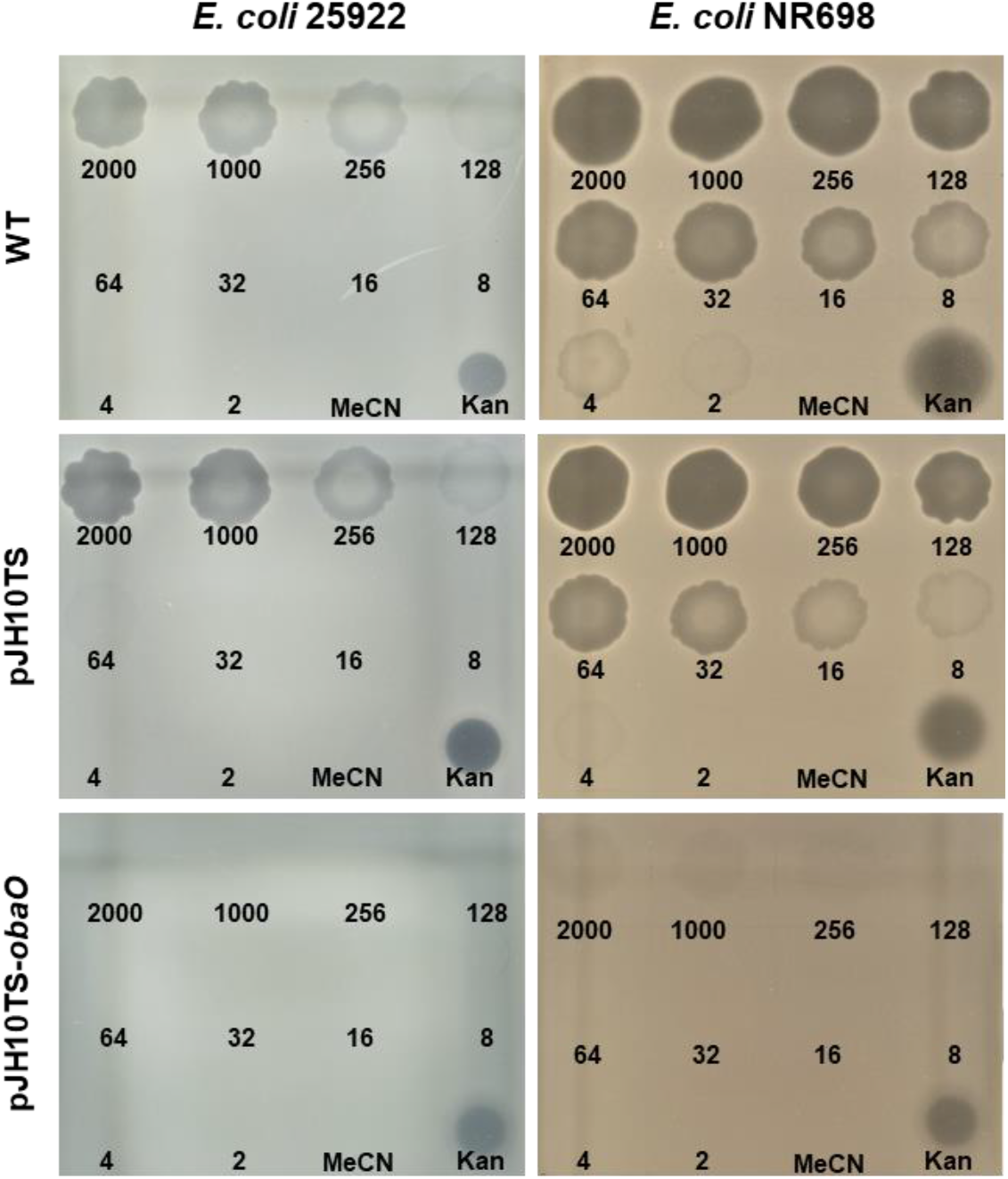
ObaO confers transferable immunity to 1-sensitive *E. coli* strains. **1** spot-on-lawn assays with clinical strain *E. coli* 25922 and permeable strain *E. coli* NR698, with zones of clearing indicating growth inhibition. Numbers indicate **1** concentrations in μg/mL, with MeCN as a negative control and kanamycin (50 μg/mL) as a positive control. ObaO confers resistance to >2000 μg/mL **1**, in contrast to the WT strains and empty vector controls. Images are representative of three biological repeats for each strain.

To determine whether simple overexpression of an additional copy of ThrRS is sufficient to provide resistance to **1**, we turned our attention to the primary housekeeping versions of ThrRS present in *P. fluorescens* ATCC 39502 and *E. coli* (EcThrRS; GenBank accession: WP_001144202.1). The Δ*obaL*Δ*obaO* strain was complemented by ectopic expression of the genes encoding either PfThrRS or EcThrRS (amplified from *E. coli* BL21 (DE3)) using the pJH10TS vector and was then grown in production medium supplemented with **2** to activate **1** biosynthesis. We observed that culture growth plateaued after 5 h and that the production of **1** was not restored in both cases (Fig. 5A). Similarly, we found that when *E. coli* bioassay strains expressing either an additional copy of their native EcThrRS or PfThrRS were challenged with **1**, there was only a modest increase in **1**-resistance for *E. coli* NR698, with a **1** MIC of 16 μg/mL (2-fold increase versus control). The MIC of **1** against *E. coli* 25922 was unchanged at 256 μg/mL (Fig. 5B, Table 1). This suggested that ObaO functions as the primary **1** resistance determinant.

**Fig. 5.**
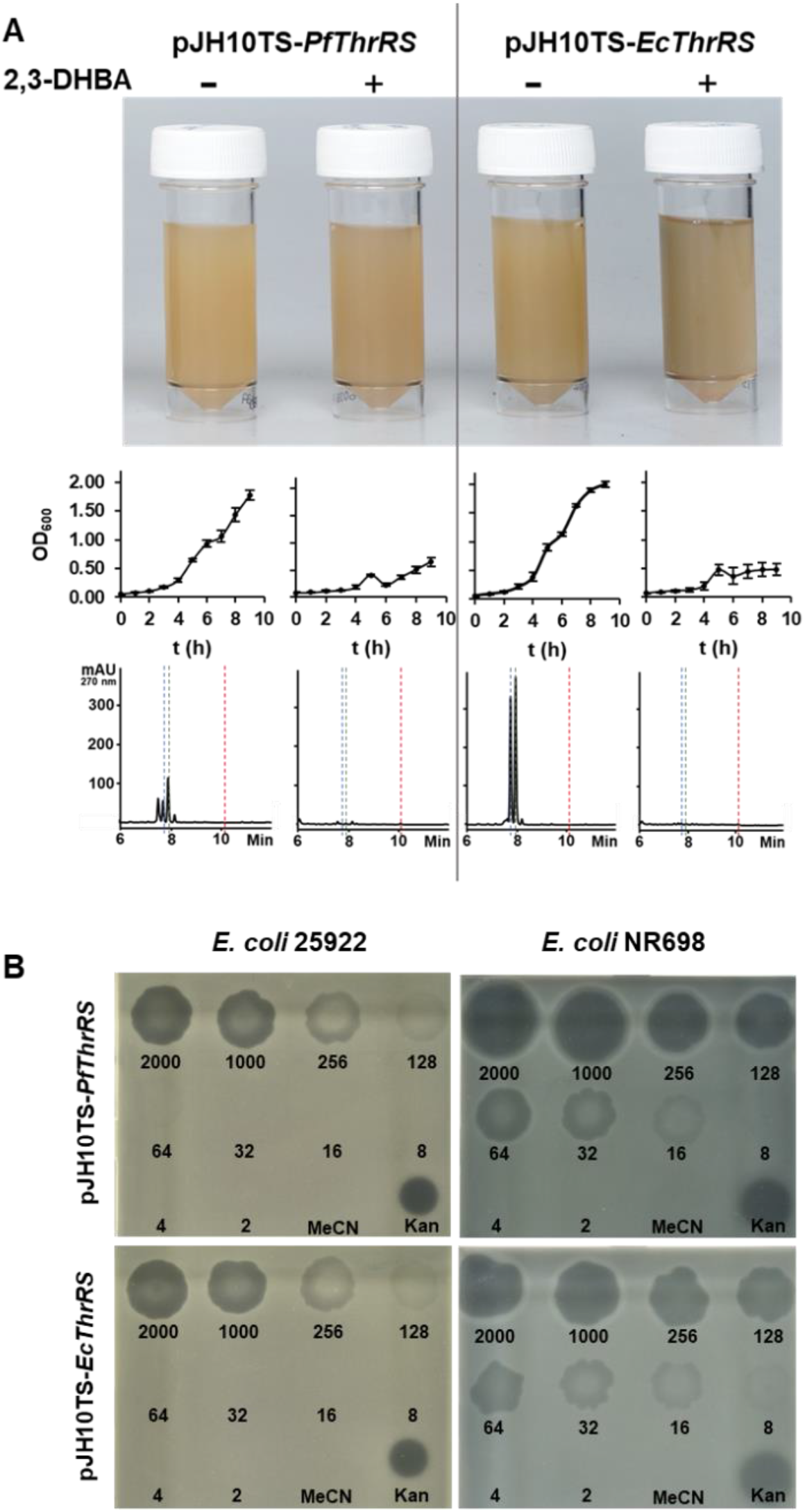
PfThrRS and EcThrRS do not confer 1-immunity. **A)** Complementation of *P. fluorescens* Δ*obaL*Δ*obaO* with ThrRS genes. Δ*obaL*Δ*obaO* was complemented with *PfThrRS* or *EcThrRS* in pJH10TS and grown ± **2** (0.2 mM). In contrast to *obaO* (Fig. 3) growth plateaued after ~5 h for *PfThrRS* and *EcThrRS* strains + **2** and **1** production was not restored. **A)** Aliquots of cultures after 14 h growth, log phase growth curves and representative HPLC chromatograms are shown, analogous to Fig. 3. Each growth curve data point is the average of three biological repeats, with bars showing the standard error and HPLC analysis was carried out on every biological repeat. **B**) **1** spot-on-lawn assays as in Fig. 4, with clinical strain *E. coli* 25922 and permeable strain *E. coli* NR698 overexpressing either PfThrRS or an additional copy of EcThrRS. In contrast to ObaO, there is no difference in the **1** MIC. Images are representative of three biological repeats for each strain.

### 1 is potent inhibitor of EcThrRS and a partial inhibitor of ObaO

To confirm that **1** is a *bona fide* inhibitor of ThrRS, we examined the formation of Thr-tRNA^Thr^ by EcThrRS in the presence of 0-5 μM **1**. The rate of aminoacylation of tRNA decreased progressively over this concentration range, with complete inhibition occurring around 1 μM (Fig. 6A). Fitting of the data to a dose response equation returned an IC_50_ of 92 ± 21 nM, confirming that **1** is a potent inhibitor of EcThrRS (Fig. 6C). Aminoacylation assays with ObaO indicated that, in the absence of **1**, ObaO exhibits activity comparable to EcThrRS. When ObaO was challenged with increasing concentrations of **1**, aminoacylation was initially decreased but not completely inhibited at the highest concentrations of **1** (Fig. 6B). In order to fit the data and derive an IC_50_, a modified dose response equation was employed that includes a term denoting residual enzyme activity at saturating inhibitor concentrations.^26^ Fitting the data to this equation gave an IC_50_ of 50 ± 25 nM and a fractional residual activity of 0.35 ± 0.07 (Fig. 6D). These data confirm that ObaO provides protection against **1** to its cellular host cell *via* a mechanism involving partial inhibition.

**Fig. 6.**
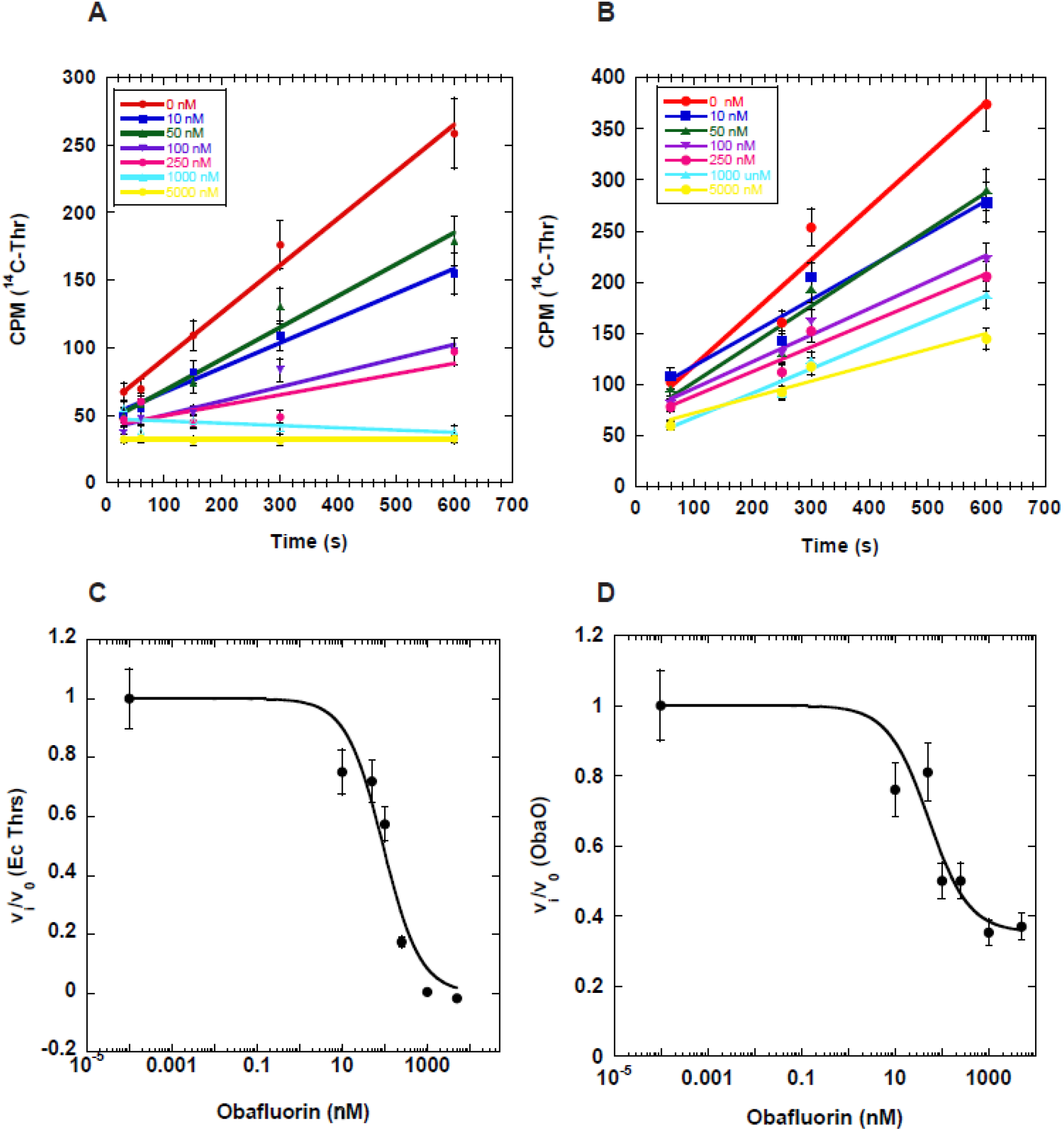
Complete and partial Inhibition of EcThrRS and ObaO, respectively, by **1**. Progress curves for EcThrRS (**A**) and ObaO (**B**) in the presence of varying (0 – 5 μM) concentrations of **1**. Enzyme (10 nM) and varying concentrations of **1** were pre-incubated for 30 min prior to adding to a reaction containing saturating substrates. Reactions (n = 3) included enzyme at 10 nM and saturating concentration of tRNA, threonine, and ATP. The progress curves were fit to a linear equation to derive initial rates. Error bars represent the standard error for each time point. Dose response curves for EcThrRS (**C**) and ObaO (**D**) were calculated from the data in A and B by plotting the fractional velocity at each of seven different inhibitor concentrations against log [**1**]. Details of the fitting routines are presented in the ESI.

## Discussion

Natural products, especially those produced by microorganisms, account for approximately 80% of all antibiotics in clinical use.^27^ Moreover, numerous other natural products have been identified with potent antibacterial activity but have failed to progress into clinical use, or even investigation, due to confounding activities such as off target effects, poor bioavailability, or a lack of broadspectrum activity. Given recent predictions that infectious disease will become the biggest killer of humans by 2050 due to the alarming increase in antimicrobial resistance,^28^ reevaluation of these previously discarded natural products is an area of great interest. By focusing on structurally unique molecules we hypothesize that new targets and modes of action might be uncovered and provide an opportunity to avoid cross resistance with existing clinical classes. One such molecule is the *β*-lactone antibiotic **1**.

Using a combination of bioinformatics, mutational analyses, bioassays, and *in vitro* enzyme assays we identified the gene product ObaO as the immunity determinant of **1**, which functions as a resistant isoform of the **1** target ThrRS. Self-immunity guided natural product target discovery has been the subject of much interest in recent years,^16,29^ however there are still relatively few examples in which a hitherto unknown target has been identified *de novo* from its self-immunity determinant.^30,31^ Our data demonstrate the utility of this approach, especially in cases such as ours, when the inherent lability of the compound in aqueous solution precludes the generation of spontaneous mutants.

aaRSs are attractive therapeutic targets, due to their essential role in translation and the potential to exploit divergences in prokaryotic and eukaryotic forms to generate selective drugs.^19,20^ aaRS-targeting natural products have been applied in both crop protection and clinical use, with the LeuRS inhibitor Agrocin TM84 used to treat crown gall disease^32^ and the IleRS inhibitor mupirocin used topically in humans to treat skin infections.^33^ A variety of mechanisms for aaRS inhibition have been reported, with compounds targeting any of the amino acid, ATP or tRNA subsites within the enzyme, as well as the tRNA substrate itself.^34^ The polyketide natural product borrelidin acts by simultaneously occupying the three catalytic subsites, along with a non-catalytic fourth subsite and is the only other aaRS inhibitor targeting the ThrRS enzyme reported thus far.^35,36^

We established that while expression of an extra copy of the native PfThrRS or the *E. coli* EcThrRS could restore some growth in the *obaO* deficient *P. fluorescens* strain, only complementation with ObaO restored WT-like growth and **1** production. The growth plateau observed for the PfThrRS and EcThrRS complemented strains may be due to L-threonine starvation resulting in the stringent response, as has been suggested previously.^37^ We also showed that expression of EcThRS and PfThrRS did not significantly alter the MIC for **1** in *E. coli* strains. These data indicate that ObaO does not provide immunity by a simple increase in levels of a functional ThrRS.

The mode of self-resistance we report here has been demonstrated for other natural products that target aaRSs. Most notably, mupirocin, for which a mupirocin-resistant IleRS, MupM, is encoded in the mupirocin BGC.^21^ Comparison of mupirocin resistant and susceptible IleRS sequences in *Thermus thermophilus* revealed that they differ by only two amino acids in the binding site.^38^ A further example is indolmycin for which *ind0* encodes a putative resistant TrpRS,^29^ however the resistance determinants have not yet been fully elucidated. Initial comparison of EcThrRS and ObaO show that they share 54% sequence identity, with conservation of all amino acid side chains that interact with tRNA, the catalytic Zn^2+^ ion and the Thr-AMP substrate intermediate (ESI Fig. 2).^39^ The basis of **1**-resistance in ObaO will be the subject of further in-depth study.

Biochemical analysis showed that whereas **1** inhibits EcThrRS completely at approximately 10x the IC_50_, ObaO retains 35% of its activity even in the presence of 100x the IC_50_ concentration. Such partial inhibition is a rarely reported phenomenon, whereby the activity of an enzyme cannot be driven to zero, even at the highest concentrations of inhibitor. This observation is often attributed to experimental artefacts, most commonly limited compound solubility at high concentrations.^26^ However, the contrasting EcThrRS and ObaO profiles (Fig. 6) with the same **1** concentration ranges, clearly exclude that possibility in this case. Previous examples of partial inhibition include the inhibition of reverse transcriptase by the non-nucleoside inhibitor tetrahydro-benzodiazepine^40^ and inhibition of tyrosinase by substituted benzaldehydes.^41^ The details of the ObaO partial inhibition mechanism remain to be fully characterized in future work. To the best of our knowledge this is the first time a partial inhibition mechanism has been implicated in the immunity to a microbial natural product.

In summary, our combined data strongly support the hypothesis that ThrRS is the molecular target of the *β*-lactone antibiotic **1** in sensitive bacteria, and that ObaO, a homologue of ThrRS, is the immunity-determinant in **1** producing bacteria. Our findings set the scene for future study on the mechanism by which **1** inhibits sensitive ThrRS, and into the resistance determinants of ObaO. In turn, this will enable rational drug design to generate improved versions of obafluorin.

## Experimental

For details regarding experimental procedures see ESI.

## Supporting information

Electronic supplementary information

## Acknowledgements

This work was supported by the Biotechnology and Biological Sciences Research Council *via* institute strategic program BBS/E/J/00PR9791 (to the John Innes Centre (JIC)), and research grant BB/P021506/1 to B.W. The contributions of P.W. and C.S.F. were supported by National Institute of General Medical Science grant GM54899-20. We thank Dr Lionel Hill and Dr Gerhard Saalbach (JIC) for excellent metabolomics support.

