## Supplementary material for "Self-immunity guided identification of threonyl-tRNA synthetase as the molecular target of obafluorin, a *β*-lactone antibiotic": Electronic supplementary information

[c] Current address: Institute of Microbiology, Eidgenössische Technische Hochschule (ETH) Zürich, Vladimir-Prelog-Weg 4, 8093 Zürich, Switzerland

#### Table of Contents

1. Materials and Methods

2. Supplementary Figures

3. Supplementary Tables

4. Gene and Protein Sequences

### 1. Materials and Methods

#### General

Reagents and chemicals were purchased from Sigma-Aldrich unless otherwise stated and used without further purification. HPLC grade solvents were purchased from Fisher Scientific. All primers, plasmids and strains are reported in Supplementary Tables 2-3. All strains were maintained on lysogeny broth (LB) medium with appropriate selection at 37 °C for *E. coli* strains or 28 °C for *Pseudomonas fluorescens* ATCC 39502 strains.

#### Construction of knockout, complementation and expression plasmids

Plasmids were constructed using standard ligation methods or using a Gibson Assembly® Cloning Kit (NEB) according to the manufacturer's instructions. Gene sequences were amplified from genomic DNA templates and vector backbones were prepared by either digestion with restriction endonucleases or PCR-amplification from plasmid DNA.

Δ*obaO* strains were generated using the suicide vector pTS1, constructed previously.[1] Primers were designed to amplify 800–1,200 bp flanking regions of the *obaO* protein coding sequence (PCS) for cloning into pTS1 between *Xba*I and *Avr*II sites, and *Avr*II and *Bmt*I sites. Flanking regions were designed to comprise 10-50 PCS codons at either end of the *obaO* gene to minimize polar effects, leaving a truncated chromosomal copy of the gene with an in-frame deletion and internal *Avr*II site cloning artefact following double homologous recombination.

For genetic complementation, the *obaO*, *E. coli* threonyl-tRNA synthetase (*EcThrRS*) and *P. fluorescens* native threonyl-tRNA synthetase (*PfThrRS*) PCSs were cloned from start to stop either as *Bmt*I-*Kpn*I fragments (*obaO* and *EcThrRS*) or as a *Nde*I-*Xba*I fragment (*PfThrRS*). These were ligated into pJH10TS[1] for introduction and ectopic expression in the Δ*obaL*Δ*obaO* strain. The *obaO* and *EcThrRS* PCSs were also cloned into pET28a(+) as *Nde*I-*Xho*I fragments, for expression of N-terminally hexahistidine tagged proteins.

#### Mutagenesis and complementation experiments.

pTS1-*obaO* knockout constructs were introduced into *P. fluorescens* ATCC 39502 strains via conjugation from *E. coli* S17-1 λpir and single-crossover mutants were selected for on LB supplemented with 50 mg/mL tetracycline ($Tc^{50}$). Positive colonies were cultured overnight in antibiotic-free medium to allow time for a second cross-over event to occur. *sacB* counter-selection could then be performed by plating culture dilutions on LB supplemented with 10% sucrose to select against retention of the pTS1 vector backbone. Colony PCR was then performed to distinguish double cross-over mutants from WT colonies, with subsequent Sanger sequencing (Eurofins Genomics) to confirm the expected deletion.

For the WT strain, all double crossover mutants retained the *obaO* gene, presumably due to selection against losing the obafluorin (**1**) resistance gene. Consistent with this hypothesis, it was possible to delete *obaO* in the previously reported Δ*obaL* strain[1], in which **1** biosynthesis is abolished. The resulting Δ*obaL*Δ*obaO* strain was subsequently complemented with pJH10TS-*obaO*, -*EcThrRS* and -*PfThrRS* constructs, or a pJH10TS empty vector control, via conjugation and positive clones were selected for on LB $Tc^{50}$. Clones were screened by colony PCR and confirmed by sequencing.

**Analysis of growth and metabolite production.**

WT and recombinant *P. fluorescens* ATCC 39502 strains were grown in **1** Production Medium (OPM) comprising: yeast extract 0.5%, D-glucose 0.5%, $MgSO_4.7H_2O$ 0.01%, and $FeSO_4$ 0.01%, dissolved in Milli-Q (Merck Millipore) filtered water. A toothpick was used to inoculate 100 mL of OPM seed culture (250 mL Erlenmeyer flask) from a single colony, with subsequent growth for 24 h at 25 °C, 300 rpm. 1 mL of this culture was used to inoculate 100 mL (500 mL Erlenmeyer flask) OPM production cultures, which were supplemented with either 2,3-dihydroxybenzoic acid (**2**) in DMSO to a final concentration of 0.2 mM or DMSO only, to give a final concentration of 0.2% DMSO in both cases.

For growth curves, production cultures were grown at 25 °C, 300 rpm and $OD_{600}$ measurements relative to an OPM blank were recorded every hour for 10 h. For metabolite analysis, after 14 h of growth 1 mL of culture broth was extracted with an equal volume of ethyl acetate by mixing at 1,400 rpm for 15 min. Samples were then centrifuged (15,682 × *g* for 15 min), and the organic phase was collected and evaporated. The resulting extract was dissolved in MeCN (250 μL) and centrifuged (15,682 × *g* for 20 min) to remove any remaining cell debris, before HPLC analyses.

**Analytical HPLC**

Samples were analysed on an Agilent 1100 system using a Gemini 3 μm NX-C18 110 Å, 150 x 4.6 mm column (Phenomenex) with a gradient elution: MeCN/0.1% (v/v) TFA ($H_2O$) gradient from 10/90 to 100/0 0–15 min, 100/0 for 15–16 min, gradient to 10/90 16–16.50 min and 10/90 for 16.50–23 min. The flow rate was 1 mL/min and the injection volume of each sample 10 μL.

Chromatograms were recorded at 270 nm, the $\lambda_{max}$ for **1**. The identity of **1** peaks were confirmed by comparison with an authentic standard. Retention times were: obafluorin 10.1

min, 4-nitrophenylethanol 7.7 min and 4-nitropenylacetate 7.9 min, the latter two compounds being shunt metabolites which are elevated in non-complemented Δ*obaL* strains.

**Antibacterial assays**

For agar diffusion bioassays, test strains were grown for 16-18 h in 5 mL LB cultures, containing appropriate selection. 500 μL of each culture was used to inoculate 50 mL LB cultures, which were incubated at 37 °C with 250 rpm until $OD_{600}$ = 0.3-0.4. Cultures were diluted 1:10 with molten soft nutrient agar (SNA), before pouring into appropriately sized petri dishes to set. Serial dilutions of **1** were prepared in MeCN and 4 μL of each dilution was applied directly onto the SNA surface. Kanamycin (50 μg/mL) was used as a positive control, and MeCN as a negative control. Plates were incubated at 25 °C for 16-18 h. The minimum inhibitory concentration (MIC) was defined as the lowest concentration of compound that inhibited resulted in a zone of inhibition. Experiments were carried out in at least triplicate for each strain.

**Protein expression and purification**

*E. coli* NiCo21(DE3) (NEB) carrying pLysS and either pET28a(+)-*EcThrRS* or pET28a(+)-*obaO* was cultivated in Terrific Broth (TB) at 28 °C and 250 rpm on a rotary shaker until $OD_{600}$ = 0.5 - 0.6. Protein expression was induced by addition of 0.1 mM IPTG and incubation continued at 18 °C and 200 rpm for 18 h. Cells were pelleted at 2,415 × *g* at 4 °C and were subsequently re-suspended in lysis buffer containing 25 mM Tris/HCl at pH 8.0 containing NaCl (300 mM), $MgCl_2$ (10 mM) and glycerol (10%). After disruption with an EmulsiFlex-B15 high pressure homogeniser (Avestin, Inc.), cells were pelleted at 26,892 × *g* at 4 °C for 30 min. The lysed supernatant was incubated with chitin resin with gentle mixing for 30 min to remove any endogenous *E. coli* metal binding proteins. Eluted sample was loaded onto a HisTrap excel (GE Healthcare) Ni-NTA column using an ÄKTA pure (GE Healthcare) system. Proteins were washed in 5 CV of lysis buffer containing 10, 20, 30 and 50 mM imidazole concentrations and then eluted with 20 CV of 250 mM imidazole and collected in in 2 mL fractions. Fractions containing protein of the expected size were determined by SDS-PAGE, pooled, diluted to remove imidazole and concentrated using Amincon columns (30 kDa MWCO). Proteins were further purified by size exclusion over a HiLoad 16/600 Superdex 200 pg column (GE Healthcare) and fractions were combined and concentrated as above. Protein samples were stored -80 °C before in vitro assays. Protein identities were confirmed by excising bands from SDS-PAGE gels, subjecting them to tryptic digest according to standard procedures adapted from Shevchenko et al.[2] and LCMS/MS analysis of peptide fragments with an Orbitrap-Fusion™ mass spectrometer (Thermo Fisher).

#### In vivo tRNA$^{Thr}$ transcription and purification

To obtain purified tRNA$^{Thr}$ for our kinetics and aminoacylation assays, tRNA$^{Thr}$ was overexpressed in *E. coli* and purified by gel electrophoresis and electroelution as described in previous works from the Francklyn Lab.[3] *E. coli* tRNA$^{Thr}$ was expressed in BL21 *E. coli* cells and purified via phenol chloroform extraction. The tRNA$^{Thr}$ was then precipitated overnight in 2.5x volume EtOH and 0.1x volume sodium acetate and subjected to centrifugation. After washing the pellet with 75% EtOH it was resuspended in 10 mM HEPES pH 6.0. This sample was then mixed with 6x blue loading dye and loaded into a large urea gel (6.5% polyacrylamide (19:1 acrylamide:bisacrylamide), 8 M urea, and 0.5 M sodium acetate pH 5.0). The gel was subject to 50 W until the dye front almost ran off the gel, at which point the gel was imaged via a UV light box and the tRNA$^{Thr}$ band was identified. This band was excised, chopped and placed into an electroeluter apparatus (Whatman/Schleicher & Schuell) overnight. After electroelution, the purified sample was again precipitated with 2.5x volume EtOH and 0.5x volume sodium acetate and then resuspended in TE pH 6.0 buffer.

#### Aminoacylation Assay

To measure the effects of **1** on ThrRS canonical tRNA charging activity, an assay modified from Ruan et al.[4] was used, where active enzyme was incubated with its necessary substrates and its activity measured $^{14}$C labeled Thr and a liquid scintillation counter. Purified ThrRS protein (10 nM) was pre-incubated with varying concentrations of **1** for 10 min. After pre-incubation, ThrRS and **1** were added to a master reaction mixture with the final concentrations of 100 mM HEPES pH 7.0, 4 mM ATP, 10 mM $MgCl_2$, 50 $\mu$M $^{14}$C labeled Thr (Moravek), and 5 $\mu$M tRNA$^{Thr}$. This mixture was then incubated for 10 min, with time points being taken at 1, 2.5, 5, and 10 min. At each time point, three 5 $\mu$L aliquots were spotted onto 5% TCA presoaked 3MM Whatman paper (Sigma-Aldrich). After letting the spots dry, the Whatman paper was washed three times with 5% TCA, and once with 95% EtOH. Whatman paper was dried and the counts on each square of paper analyzed with a liquid scintillation counter using Hydrofluor Liquid Scintillation Fluid (National Diagnostics). To calculate $IC_{50}$, fractional initial velocities (velocity of inhibited reaction/velocity in the absence of inhibitor) were plotted against the log of **1** concentration and then fitted to equation 1.[5]

$$\frac{v_i}{v_o} = \frac{1}{1 + \frac{[I]}{IC_{50}}} \quad (1)$$

For ObaO, a modified version of the equation was used to take into account the observed partial inhibition:

$$y = \frac{y_{max} - y_{min}}{1 + \frac{[I]}{IC_{50}}} y_{min}$$

(2)

#### 2. Supplementary Figures

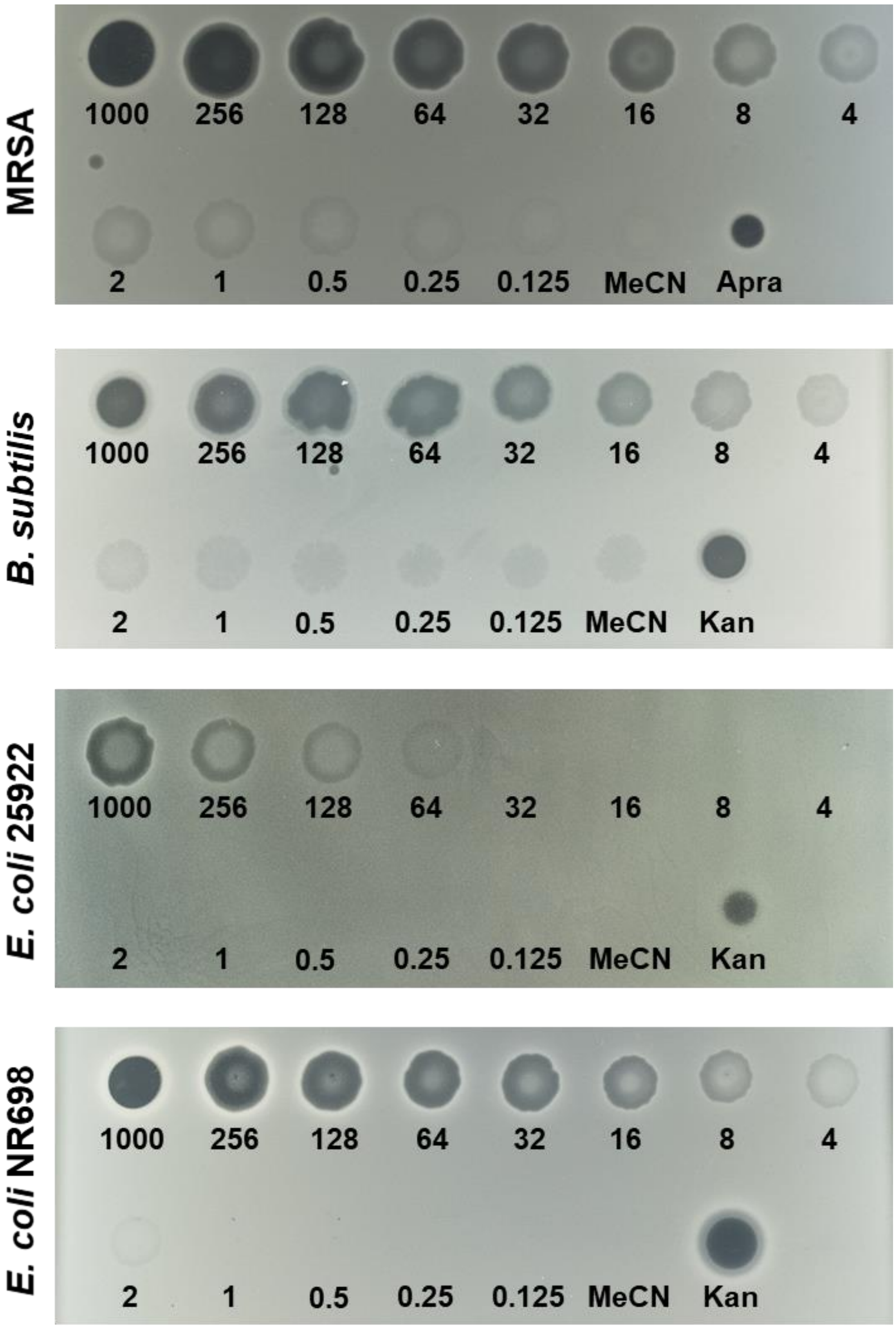


**Supplementary Fig. 1 Obafluorin is active against Gram-positive and Gram-negative bacterial strains.** Spot-on-lawn assays with *Bacillus subtilis*, methicillin resistant *Staphylococcus aureus* (MRSA), *E. coli* 25922 and *E. coli* NR698, with zones of clearing indicating growth inhibition. Numbers indicate **1** concentrations in μg/mL, with MeCN as a negative control and either apramycin or kanamycin (50 μg/mL) as a positive control. Images are representative of at least three biological repeats for each strain.

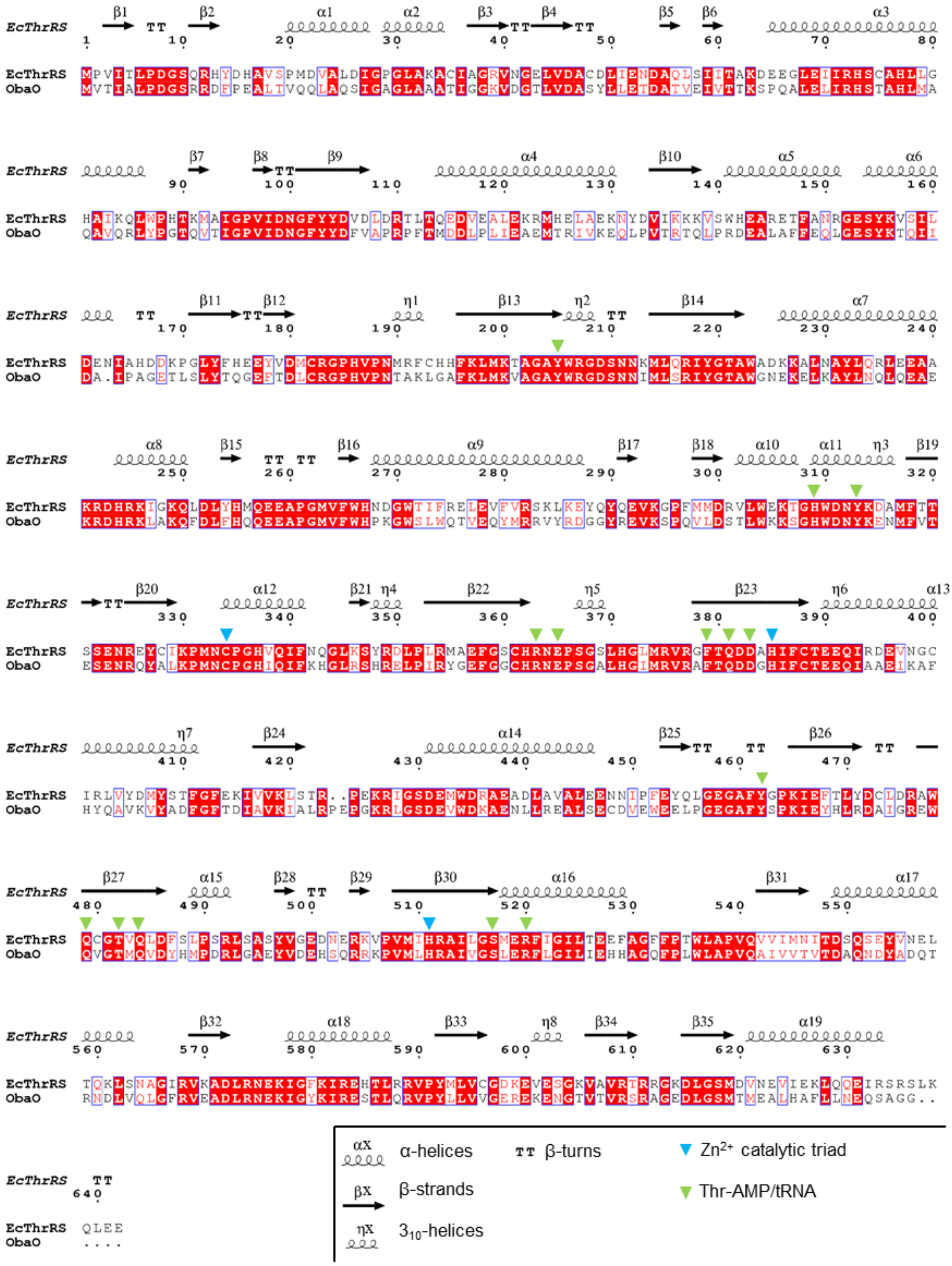


**Supplementary Fig. 2 Conservation of EcThrRS catalytic residues in ObaO.** EcThrRS side chains that interact with either the Thr-AMP and tRNA substrates or the catalytic $Zn^{2+}$ ion[6] are indicated in green and blue, respectively. The secondary structural motifs of EcThrRS (PDB ID 1QF6) are shown according to the key in the bottom right. The sequence alignment was generated using Clustal Omega[7] and displayed with ESPript.[8]

#### 3. Supplementary Tables

##### Supplementary Table 1: Obafluorin BGC homologues

| Strain | NCBI Reference Sequence | Sequence coordinates(bp) |
|---|---|---|
| *Pseudomonas fluorescens* ATCC 39502 | KX931446.1 | 1..19,561 |
| *Pseudomonas fluorescens* PfR 37 | NZ_KZ477995.1 | 70,524..49,501 |
| *Pseudomonas orientalis* F9 | NZ_CP018049.1 | 4,685,939..4,706,957 |
| *Pseudomonas orientalis* L1-3-08 | NZ_CP027724.1 | 4,668,946..4,689,253 |
| *Pseudomonas* sp. 286 | NZ_UTBO01000036.1 | 61,266..40,209* |
| *Pseudomonas* sp. 34 E 7 | NZ_CVTX01000156.1 | 76,609..97,226 |
| *Pseudomonas* sp. 37 R 15 | NZ_CVTV01000010.1 | 71,693..52,583* |
| *Pseudomonas* sp. Irchel s3a18 | NZ_FYDV01000019.1 | 44,071..22,247* |
| *Burkholderia diffusa* INT-BP16 | NZ_LOUS01000041.1 | 81,817..59,914* |
| *Burkholderia diffusa* MSMB583 | NZ_LPKA01000054.1 | 19,042..40,939 |
| *Burkholderia diffusa* MSMB 1060 | NZ_LOYB01000006.1 | 286,604..308,518 |
| *Burkholderia diffusa* RF8-non_BP2 | NZ_LOTC01000033.1 | 80,074.58,171* |
| *Burkholderia multivorans* AU21015 | NZ_FKJW01000005.1 | 254,862..276,758 |
| *Burkholderia multivorans* LMG29306 | NZ_FKJV01000003.1 | 265,699..287,595 |
| *Burkholderia multivorans* LMG29311 | NZ_FKJW01000005.1 | 320,892..342,788 |
| *Burkholderia* sp. Bp8998 | NZ_QTQF01000003.1 | 220,477..242,401 |
| *Burkholderia stagnalis* Bp8965 | NZ_QTPN01000035.1 | 71,071..50,304* |
| *Burkholderia stagnalis* Bp8966 | NZ_QTPM01000036.1 | 70,950..50,183* |
| *Burkholderia stagnalis* Bp8971 | NZ_QTPL01000033.1 | 71,071..50,304* |
| *Burkholderia stagnalis* Bp8975 | NZ_QTPK01000039.1 | 70,950..50,183* |
| *Burkholderia stagnalis* Bp8976 | NZ_QTPJ01000036.1 | 71,080..50,313* |
| *Burkholderia stagnalis* Bp9121 | NZ_QTOP01000034.1 | 71,071..50,304* |
| *Burkholderia stagnalis* Bp9122 | NZ_QTOO01000036.1 | 70,950..50,183* |
| *Burkholderia stagnalis* Bp9123 | NZ_QTON01000038.1 | 70,950..50,183* |
| *Burkholderia stagnalis* Bp9124 | NZ_QTOM01000037.1 | 70,950..50,183* |
| *Burkholderia stagnalis* Bp9151 | NZ_QTOI01000037.1 | 70,950..50,183* |
| *Burkholderia stagnalis* MSMB653 | NZ_LPKN01000088.1 | 70,998..50,231* |
| *Burkholderia stagnalis* MSMB654 | NZ_LPKO01000021.1 | 70,998..50,231* |
| *Burkholderia stagnalis* MSMB749 | NZ_LPLJ01000056.1 | 71,035..50,268* |
| *Burkholderia stagnalis* MSMB756 | NZ_LPLM01000075.1 | 70,998..50,231* |
| *Burkholderia stagnalis* MSMB757 | NZ_LPLN01000100.1 | 42,728..63,495 |
| *Burkholderia stagnalis* MSMB808 | NZ_LPME01000012.1 | 70,998..50,231* |
| *Burkholderia stagnalis* MSMB809 | NZ_LPMF01000025.1 | 70,998..50,231* |
| *Burkholderia stagnalis* MSMB810 | NZ_LPMG01000095.1 | 70,998..50,231* |
| *Burkholderia stagnalis* MSMB835 | NZ_LPMM01000075.1 | 70,998..50,231* |
| *Burkholderia stagnalis* MSMB836 | NZ_LPMN01000027.1 | 70,998..50,231* |
| *Burkholderia stagnalis* MSMB837 | NZ_LPMO01000030.1 | 70,998..50,231* |
| *Burkholderia stagnalis* MSMB845 | NZ_LPMR01000053.1 | 71,035..50,268* |
| *Burkholderia stagnalis* MSMB1511 | NZ_LPMY01000093.1 | 70,998..50,231* |
| *Burkholderia stagnalis* MSMB1521 | NZ_LPGD01000044.1 | 70,998..50,231* |
| *Burkholderia stagnalis* MSMB1943 | NZ_LPGZ01000073.1 | 70,998..50,231* |
| *Burkholderia stagnalis* MSMB1956 | NZ_LPHA01000089.1 | 70,998..50,231* |
| *Burkholderia stagnalis* MSMB1960 | NZ_LPHB01000055.1 | 70,998..50,231* |
| *Burkholderia stagnalis* MSMB2168 | NZ_LPIH01000025.1 | 70,998..50,231* |
| *Burkholderia stagnalis* MSMB2169 | NZ_LPII01000020.1 | 70,998..50,231* |
| *Burkholderia territorii* RF6-non_BP1 | NZ_LOSY01000044.1 | 265,090..286,999 |
| *Burkholderia territorii* RF7-non_BP1 | NZ_LOTB01000027.1 | 265,090..286,999 |
| *Chitiniphilus shinanonensis* DSM 23277 | NZ_KB895358.1 | 15,542..36,162 |

#### Supplementary Table 2: Strains and plasmids used in this work.

| Plasmid/Strain | Description | Reference |
|---|---|---|
| **Plasmid** | | |
| pTS1 | pME3087 modified with *sacB* counter-selection and an expanded multiple cloning site | 1 |
| pJH10TS | pJH10 modified with an expanded cloning site | 1 |
| pET28a(+) | Expression vector; Kan$^R$, the transcription of the cloned gene is driven by the T7 RNA polymerase and controlled by the LacI repressor, *ColE1* replicon | Novagen |
| pLysS | Vector for basal expression from the T7 promoter by producing T7 lysozyme; p15A replicon, Cm$^R$ | Novagen |
| **Strain** | | |
| *E. coli* DH5α | Cloning strain; F$^-$ *endA1 glnV44 thi-1 recA1 relA1 gyrA96 deoR nupG* ϕ80*dlac*Δ*(lacZ)M15* Δ(*lacIZYA-argF*)U169 *hsdR*17(r$_K^-$ m$_K^+$) λ– | Lab stock |
| *E. coli* S17-1 λ(pir) | Donor strain for conjugation between *E. coli* and *P. fluorescens* ATCC 39502; *recA thi pro hsd*(R$^-$ M$^+$)RP4: 2-Tc::Mu- Km::Tn7 λpir SM$^R$ Tp$^R$ | Lab stock |
| *E. coli* BL21(DE3) | Expression strain; *fhuA2 [lon] ompT gal (λ DE3) [dcm]* Δ*hsdS* λ *DE3 =* λ *sBam*HIo Δ*Eco*RI-*B int::(lacI::PlacUV5::T7 gene1) i21* Δ*nin5* | Lab stock |
| *E. coli* NiCo21 (DE3) | Expression strain; *can::CBD*, *fhuA2*, *[lon] ompT*, *gal* (λ DE3) *[dcm] arnA::CBD*, *slyD::CBD*, *glmS6Ala*, Δ*hsdS* λ DE3 = λ *sBamHIo* Δ*EcoRI-B int::(lacI::PlacUV5::T7 gene1) i21*, Δ*nin5*. | NEB |
| *E. coli* ATCC 25922 | Bioassay strain; WT | ATCC, USA |
| *E. coli* NR698 | Bioassay strain; MC4100 (F$^-$ *araD139* Δ(*argF-lac*)*U169*, *rpsL150*, *relA1*, *flbB5301*, *deoC1*, *ptsF25*, *rbsR*), *imp4213* | 9 |
| *B. subtilis* | Laboratory strain provided by the Handelsman lab | 10 |
| *MRSA* | Methicillin-resistant *S. aureus*; clinical isolate provided by Dr Justin O'Grady (UEA Medical School) | 10 |
| *P. fluorescens* ATCC 39502 | WT obafluorin producing strain | ATCC, USA |
| *P. fluorescens* Δ*obaL* | *P. fluorescens* ATCC 39502 with an in-frame truncation in the *obaL* gene | 1 |
| *P. fluorescens* Δ*obaL* Δ*obaO* | ATCC 39502 with an in-frame truncation in the *obaL* and *obaO* genes | This work |
| *P. fluorescens* Δ*obaL* Δ*obaO* pJH10TS | Δ*obaL* Δ*obaO* strain carrying the empty pJH10TS plasmid as a negative control | This work |
| *P. fluorescens* Δ*obaL* Δ*obaO* pJH10TS-*obaO* | Δ*obaL* Δ*obaO* strain complemented with *obaO* | This work |
| *P. fluorescens* Δ*obaL* Δ*obaO* pJH10TS-*EcThrRS* | Δ*obaL* Δ*obaO* strain complemented with *EcThrRS* | This work |
| *P. fluorescens* Δ*obaL* Δ*obaO* pJH10TS-*PfThrRS* | Δ*obaL* Δ*obaO* strain complemented with an additional copy of *PfThrRS* | This work |
| *E. coli* ATCC 25922 pJH10TS | 25922 carrying the pJH10TS plasmid as a negative control | This work |
| *E. coli* ATCC 25922 pJH10TS-*obaO* | 25922 carrying pJH10TS-*obaO* | This work |
| *E. coli* ATCC 25922 pJH10TS-*EcThrRS* | 25922 carrying pJH10TS-*EcThRS* | This work |
| *E. coli* ATCC 25922 pJH10TS-*PfThrRS* | 25922 carrying pJH10TS-*PfThrRS* | This work |
| *E. coli* NR698 pJH10TS | NR698 carrying the pJH10TS plasmid as a negative control | This work |
| *E. coli* NR698 pJH10TS-*obaO* | NR698 carrying pJH10TS-*obaO* | This work |
| *E. coli* NR698 pJH10TS-*EcThrRS* | NR698 carrying pJH10TS-*EcThRS* | This work |
| *E. coli* NR698 pJH10TS-*PfThrRS* | NR698 carrying pJH10TS-*PfThrRS* | This work |
| *E. coli* NiCo21 (DE3) pLysS pET28a(+)-*EcThrRS* | NiCo21 (DE3):pLysS carrying the pET28(+)-*EcThrRS* plasmid, for production of the His$_6$-EcThrRS protein | This work |
| *E. coli* NiCo21 (DE3) pLysS pET28a(+)-*obaO* | NiCo21 (DE3):pLysS carrying the pET28(+)-*obaO* plasmid, for production of the His$_6$-ObaO protein | This work |

**Supplementary Table 3: Oligonucleotides used in this work.** Restriction sites are indicated in bold and start codons are underlined.

| Oligonucleotide | Sequence 5' - 3' | Description |
|---|---|---|
| *obaO* KOF1 FW | CGTCGG**TCTAGA**CGCATTACGAGTTCGTCTTCCG | Each pair of KOF (KnockOut Fragment) primers is designed to amplify one upstream (1) and one downstream (2) product, overlapping the N- and C-termini of each *oba* gene coding sequence. KOF1 is cloned as an *Xba*I-*Avr*II fragment, and KOF2 as an *Avr*II-*Bmt*I, into the suicide vector pTS1. |
| *obaO* KOF1 RV | CGTCGG**CCTAGG**CTGTTGCACAGTCAAAGCTTCTGG | |
| *obaO* KOF2 FW | GCTAGT**TCTAGA**CGTCGGCCTAGGGACTTGGGAAGCATGACGATGG | |
| *obaO* KOF2 RV | GCTAGT**AAGCTT**CGTCGGGCTAGCCTGCAACTTAAGCAGGTACCCG | |
| pJH10TS-*obaO* FW | CCTGAA**GCTAGC**ATGGTCACTATCGCTCTACCGG | Each pair of pJH10TS primers is designed to clone the entire PCS of each ThrRS gene as either a *Bmt*I-*Kpn*I or *Nde*I-*Xba*I fragment into pJH10TS. |
| pJH10TS-*obaO* RV | GAGTCC**GGTACC**TCAGCCGCCTGCTGATTGC | |
| pJH10TS-*EcThrRS* FW | CCTGAA**GCTAGC**ATGCCTGTTATAACTCTTCCTGATGGC | |
| pJH10TS-*EcThrRS* RV | GAGTCC**GGTACC**TTATTCCTCCAATTGTTTAAGACTGCGGC | |
| pJH10TS-*PfThrRS* FW | TAGCACCTCTCGAGGCAT**CATATG**CCAACTATTACTCTACCC | |
| pJH10TS-*PfThrRS* RV | CACGCTCTCCAGCG**AGATCT**TTACTCCGAATCTGGGCG | |
| pET28a(+)-*obaO* FW | TGGTGCCGCGCGGCAGC**CATATG**GTCACTATCGCTCTACCG | Each pair of pET28a(+) primers is designed to clone the entire PCS of each ThrRS gene as an *Nde*I-*Xho*I fragment into pET28a(+). |
| pET28a(+)-*obaO* RV | CAGTGGTGGTGGTGGTGGTG**CTCGAG**TCAGCCGCCTGCTGATTG | |
| pET28a(+)-*EcThrRS* FW | TGGTGCCGCGCGGCAGC**CATATG**CCTGTTATAACTCTTC | |
| pET28a(+)-*EcThrRS* RV | CAGTGGTGGTGGTGGTGGTG**CTCGAG**TTATTCCTCCAATTGTTTAAGAC | |

#### 4. Gene and Protein Sequences

##### *obaO* (GenBank Accession KX931446.2)

ATGGTCACTATCGCTCTACCGGACGGCAGTCGCAGAGATTTTCCAGAAGCTTTGACTGTGCAACAGCT
GGCTCAATCCATTGGCGCAGGCCTCGCTGCCGCGACGATTGGCGGCAAGGTCGACGGCACGCTGGTCG
ATGCCAGCTACCTCCTGGAAACAGACGCTACCGTAGAGATTGTCACGACCAAAAGCCCGCAAGCGCTG
GAGCTGATACGCCATTCGACGGCCCACTTGATGGCACAGGCGGTTCAGCGCCTGTACCCCGGCACGCA
AGTAACGATAGGCCCCGTTATTGATAATGGCTTCTATTACGACTTCGTGGCACCTCGTCCTTTCACAA
TGGACGATCTTCCGCTCATCGAAGCCGAAATGACCAGGATCGTCAAAGAACAACTACCGGTTACCCGT
ACGCAATTACCACGGGATGAAGCACTGGCGTTTTTTGAACAACTGGGCGAAAGCTACAAGACGCAAAT
CATCGACGCCATACCCGCCGGGGAAACGCTCTCACTTTATACCCAGGGCGAGTTCACCGACCTGTGCC
GCGGGCCGCATGTGCCCAACACAGCGAAGCTTGGCGCCTTCAAACTGATGAAAGTGGCTGGCGCCTAC
TGGCGCGGCGACTCCAACAACATCATGCTCAGCCGTATCTACGGCACTGCCTGGGGCAACGAAAAAGA
GCTCAAGGCCTACTTGAATCAACTTCAAGAAGCTGAAAAGCGTGATCACCGCAAACTGGCAAAACAGT
TCGACTTGTTTCACCAACAAGAAGAAGCGCCAGGCATGGTCTTTTGGCACCCCAAGGGCTGGTCGTTG
TGGCAGACGGTTGAACAGTACATGCGCCGGGTTTACCGCGATGGCGGCTACCGTGAAGTCAAGTCGCC
GCAAGTACTCGACAGCACCCTGTGGAAGAAATCAGGGCACTGGGACAACTACAAAGAAAACATGTTTG
TCACTGAGTCCGAAAACCGCCAGTACGCACTCAAGCCGATGAACTGCCCCGGCCACATCCAAATCTTC
AAGCATGGCCTGCGCAGCCACCGTGAACTGCCGATCCGTTACGGCGAGTTCGGCGGCTGCCACCGCAA
CGAGCCTTCGGGCGCTTTGCACGGCATTATGCGGGTGCGCGCATTTACCCAGGATGACGGGCACATTT
TCTGCACCGAAGAGCAAATCGCGGCCGAGATCAAGGCGTTCCATTACCAGGCAGTCAAGGTTTATGCC
GACTTCGGCTTCACTGACATTGCGGTAAAAATCGCACTGCGTCCTGAGCCAGGCAAACGCCTGGGCAG
CGACGAAGTGTGGGACAAGGCCGAAAACCTGTTGCGTGAAGCCCTGTCCGAGTGTGATGTGGAGTGGG
AAGAACTGCCGGGCGAAGGCGCGTTCTACAGCCCCAAGATCGAGTACCACCTGCGTGACGCCATCGGG
CGTGAATGGCAGGTAGGCACGATGCAGGTCGACTATCACATGCCGGATCGCCTGGGCGCCGAATACGT
CGATGAGCATTCGCAACGACGCAAGCCGGTCATGCTGCACCGCGCAATCGTGGGCTCCCTGGAGCGGT
TCCTTGGCATTCTCATCGAACACCATGCCGGTCAGTTCCCGCTCTGGCTTGCGCCGGTGCAGGCCATC
GTCGTGACGGTCACTGATGCACAGAACGATTATGCCGACCAGACACGCAATGACCTGGTCCAACTTGG
GTTCAGGGTGGAAGCTGATCTGCGCAATGAGAAAATCGGCTACAAGATTCGTGAAAGCACTCTTCAAC
GCGTGCCTTACTTGCTCGTGGTCGGTGAGCGCGAAAAAGAAAACGGTACTGTCACCGTGCGCTCGCGC
GCAGGCGAAGACTTGGGAAGCATGACGATGGAAGCGCTGCACGCCTTCCTGTTGAACGAGCAATCAGC
AGGCGGCTGA

##### ObaO

MVTIALPDGSRRDFPEALTVQQLAQSIGAGLAAATIGGKVDGTLVDASYLLETDATVEIVTTKSPQAL
ELIRHSTAHLMAQAVQRLYPGTQVTIGPVIDNGFYYDFVAPRPFTMDDLPLIEAEMTRIVKEQLPVTR
TQLPRDEALAFFEQLGESYKTQIIDAIPAGETLSLYTQGEFTDLCRGPHVPNTAKLGAFKLMKVAGAY
WRGDSNNIMLSRIYGTAWGNEKELKAYLNQLQEAEKRDHRKLAKQFDLFHQQEEAPGMVFWHPKGWSL
WQTVEQYMRRVYRDGGYREVKSPQVLDSTLWKKSGHWDNYKENMFVTESENRQYALKPMNCPGHIQIF
KHGLRSHRELPIRYGEFGGCHRNEPSGALHGIMRVRAFTQDDGHIFCTEEQIAAEIKAFHYQAVKVYA
DFGFTDIAVKIALRPEPGKRLGSDEVWDKAENLLREALSECDVEWEELPGEGAFYSPKIEYHLRDAIG
REWQVGTMQVDYHMPDRLGAEYVDEHSQRRKPVMLHRAIVGSLERFLGILIEHHAGQFPLWLAPVQAI
VVTVTDAQNDYADQTRNDLVQLGFRVEADLRNEKIGYKIRESTLQRVPYLLVVGEREKENGTVTVRSR
AGEDLGSMTMEALHAFLLNEQSAGG

##### *EcThrRS* (GenBank Accession WP_001144202.1)

ATGCCTGTTATAACTCTTCCTGATGGCAGCCAACGCCATTACGATCACGCTGTAAGCCCCATGGATGT
TGCGCTGGACATTGGTCCAGGTCTGGCGAAAGCCTGTATCGCAGGGCGCGTTAATGGCGAACTGGTTG
ATGCTTGCGATCTGATTGAAAACGACGCACAACTGTCGATCATTACCGCCAAAGACGAAGAAGGTCTG
GAGATCATTCGTCACTCCTGTGCGCACCTGTTAGGGCACGCGATTAAACAACTTTGGCCGCATACCAA
AATGGCAATCGGCCCGGTTATTGACAACGGTTTTTATTACGACGTTGATCTTGACCGCACGTTAACCC
AGGAAGATGTCGAAGCACTCGAGAAGCGGATGCATGAGCTTGCTGAGAAAAACTACGACGTCATTAAG
AAGAAAGTCAGCTGGCACGAAGCGCGTGAAACTTTCGCCAACCGTGGGGAGAGCTACAAAGTCTCCAT
TCTTGACGAAAACATCGCCCATGATGACAAGCCAGGTCTGTACTTCCATGAAGAATATGTCGATATGT
GCCGCGGTCCGCACGTACCGAACATGCGTTTCTGCCATCATTTCAAACTAATGAAAACGGCAGGGGCT
TACTGGCGTGGCGACAGCAACAACAAAATGTTGCAACGTATTTACGGTACGGCGTGGGCAGACAAAAA
AGCACTTAACGCTTACCTGCAGCGCCTGGAAGAAGCCGCGAAACGCGACCACCGTAAAATCGGTAAAC
AGCTCGACCTGTACCATATGCAGGAAGAAGCGCCGGGTATGGTATTCTGGCACAACGACGGCTGGACC
ATCTTCCGTGAACTGGAAGTGTTTGTTCGTTCTAAACTGAAAGAGTACCAGTATCAGGAAGTTAAAGG
TCCGTTCATGATGGACCGTGTCCTGTGGGAAAAAACCGGTCACTGGGACAACTACAAAGATGCAATGT
TCACCACATCTTCTGAGAACCGTGAATACTGCATTAAGCCGATGAACTGCCCGGGTCACGTACAAATT
TTCAACCAGGGGCTGAAGTCTTATCGCGATCTGCCGCTGCGTATGGCCGAGTTTGGTAGCTGCCACCG
TAACGAGCCGTCAGGTTCGCTGCATGGCCTGATGCGCGTGCGTGGATTTACCCAGGATGACGCGCATA
TCTTCTGTACTGAAGAACAAATTCGCGATGAAGTTAACGGATGTATCCGTTTAGTCTATGATATGTAC
AGCACTTTTGGCTTCGAGAAGATCGTCGTCAAACTCTCCACTCGTCCTGAAAAACGTATTGGCAGCGA
CGAAATGTGGGATCGTGCTGAGGCGGACCTGGCGGTTGCGCTGGAAGAAAACAACATCCCGTTTGAAT
ATCAACTGGGTGAAGGCGCTTTCTACGGTCCGAAAATTGAATTTACCCTGTATGACTGCCTCGATCGT
GCATGGCAGTGCGGTACAGTACAGCTGGACTTCTCTTTGCCGTCTCGTCTGAGCGCTTCTTATGTAGG
CGAAGACAATGAACGTAAAGTACCGGTAATGATTCACCGCGCAATTCTGGGGTCGATGGAACGTTTCA
TCGGTATCCTGACCGAAGAGTTCGCTGGTTTCTTCCCGACCTGGCTTGCGCCGGTTCAGGTTGTTATC
ATGAATATTACCGATTCACAGTCTGAATACGTTAACGAATTGACGCAAAAACTATCAAATGCGGGCAT
TCGTGTTAAAGCAGACTTGAGAAATGAGAAGATTGGCTTTAAAATCCGCGAGCACACTTTGCGTCGCG
TCCCATATATGCTGGTCTGTGGTGATAAAGAGGTGGAATCAGGCAAAGTTGCCGTTCGCACCCGCCGT
GGTAAAGACCTGGGAAGCATGGACGTAAATGAAGTGATCGAGAAGCTGCAACAAGAGATTCGCAGCCG
CAGTCTTAAACAATTGGAGGAATAA

##### EcThrRS

MPVITLPDGSQRHYDHAVSPMDVALDIGPGLAKACIAGRVNGELVDACDLIENDAQLSIITAKDEEGL
EIIRHSCAHLLGHAIKQLWPHTKMAIGPVIDNGFYYDVDLDRTLTQEDVEALEKRMHELAEKNYDVIK
KKVSWHEARETFANRGESYKVSILDENIAHDDKPGLYFHEEYVDMCRGPHVPNMRFCHHFKLMKTAGA
YWRGDSNNKMLQRIYGTAWADKKALNAYLQRLEEAAKRDHRKIGKQLDLYHMQEEAPGMVFWHNDGWT
IFRELEVFVRSKLKEYQYQEVKGPFMMDRVLWEKTGHWDNYKDAMFTTSSENREYCIKPMNCPGHVQI
FNQGLKSYRDLPLRMAEFGSCHRNEPSGSLHGLMRVRGFTQDDAHIFCTEEQIRDEVNGCIRLVYDMY
STFGFEKIVVKLSTRPEKRIGSDEMWDRAEADLAVALEENNIPFEYQLGEGAFYGPKIEFTLYDCLDR
AWQCGTVQLDFSLPSRLSASYVGEDNERKVPVMIHRAILGSMERFIGILTEEFAGFFPTWLAPVQVVI
MNITDSQSEYVNELTQKLSNAGIRVKADLRNEKIGFKIREHTLRRVPYMLVCGDKEVESGKVAVRTRR
GKDLGSMDVNEVIEKLQQEIRSRSLKQLEE

***PfThrRS***

ATGCCAACTATTACTCTACCCGACGGCAGTCAACGTTCATTCGATCATCCGGTTTCCGTAGCCGAGGT
CGCCGCATCCATTGGTGCCGGCCTGGCCAAGGCCACCGTGGCCGGCAAGGTCGATGGCCAGCTGGTCG
ATGCCAGTGACCTGATCACCTCCGATGCCAGCCTGCAGATCATCACGCCCAAGGATCAAGAGGGGCTC
GAGATTATTCGCCACTCTTGCGCGCACCTGATTGGCCATGCGGTCAAGCAGCTGTACCCAACCGCCAA
GATGGTGATCGGCCCGGTAATCGAAGAAGGCTTCTATTACGACATCGCCTATGAGCGTCCTTTCACTC
CGGACGACCTGGCGGCCATCGAGCAGCGCATGCACGCGCTGATCGAAAAAGATTACGACGTGATCAAG
AAGGTCACCCCGCGCGCCGAAGTGATCGACGTGTTTACCGCCCGTGGCGAAGACTACAAGCTGCGCCT
GGTGGAAGACATGCCGGACGAGCAGGCCATGGGTCTGTACTACCACGAAGAATACGTGGACATGTGCC
GTGGCCCGCACGTGCCGAACACGCGCTTTCTCAAGTCGTTCAAGCTGACCAAGTTGTCCGGTGCCTAC
TGGCGCGGCGACGCAAAGAACGAGCAACTGCAACGTATCTACGGCACCGCCTGGGCCGACAAGAAGCA
GCTGGCCGCCTATATCCAGCGCATCGAAGAAGCCGAGAAACGCGACCACCGCAAGATCGGCAAGCGCC
TGAACCTGTTCCATCTGCAGGAAGAAGCGCCGGGCATGGTGTTCTGGCATCCGAACGGCTGGACCCTG
TACCAGGTGCTCGAGCAGTACATGCGCAAGGTTCAGCGCGAAAACGGCTACCTCGAGATCAAGACCCC
GCAGGTCGTCGATCGCAGCCTGTGGGAGAAGTCCGGGCACTGGGCCAACTACGCCGACAATATGTTCA
CCACCCAGTCGGAAAACCGCGACTACGCTATCAAGCCGATGAACTGCCCTTGCCATGTGCAGGTGTTC
AATCAAGGCCTGAAGAGCTACCGCGAGTTGCCGATGCGTCTGGCCGAGTTCGGTGCCTGCCACCGCAA
CGAACCATCGGGTGCGCTGCACGGCATCATGCGCGTGCGCGGCTTTACCCAGGACGACGCCCATATTT
TCTGCACCGAAGAGCAGATGCAGGCTGAATCGGCCGCGTTCATCAAGCTGACCATGGACGTTTACCGC
GATTTCGGCTTCACCGAAGTCGAGATGAAACTGTCCACTCGTCCGGAAAAACGCGTCGGCTCCGACGA
GCTCTGGGATCGCGCCGAAGCAGCACTGGCCGCAGCACTCGACAGCGCGGGCCTTGCGTACGACTTGC
AGCCGGGCGAGGGCGCGTTCTACGGTCCCAAGATCGAGTTCTCGCTGAAAGATTGCCTTGGCCGTGTC
TGGCAGTGTGGTACCCTGCAGCTCGATTTTAACCTGCCGATCCGTCTGGGAGCCGAATACGTCTGCGA
AGACAACAGTCGTAAACACCCGGTTATGCTGCACCGGGCGATCCTCGGCTCGTTCGAACGGTTCGTCG
GCATCCTGATCGAGCACTACGAGGGCGCGTTCCCGGCGTGGCTGGCTCCGACCCAGGCAGTGATCATG
AATATCACTGATAAACAGGCAGATTTTGCCGCTGAAGTTGAAAAAACACTCAACGAAAGCGGATTTCG
TGCCAAGTCCGACTTGAGAAATGAAAAGATCGGCTTTAAAATCCGTGAGCATACTTTGCTCAAGGTTC
CCTATCTTTTGGTTATCGGAGATCGGGAAGTCGAGATGCAGACTGTCGCTGTGCGTACTCGTGAAGGT
GCTGACCTGGGCTCGATGCCCGTCGCCCAGTTCGCTGAGTTCCTCGCGCAAGCGGTTTCCCGGCGTGG
TCGCCCAGATTCGGAGTAA

**PfThrRS**

MPTITLPDGSQRSFDHPVSVAEVAASIGAGLAKATVAGKVDGQLVDASDLITSDASLQIITPKDQEGL
EIIRHSCAHLIGHAVKQLYPTAKMVIGPVIEEGFYYDIAYERPFTPDDLAAIEQRMHALIEKDYDVIK
KVTPRAEVIDVFTARGEDYKLRLVEDMPDEQAMGLYYHEEYVDMCRGPHVPNTRFLKSFKLTKLSGAY
WRGDAKNEQLQRIYGTAWADKKQLAAYIQRIEEAEKRDHRKIGKRLNLFHLQEEAPGMVFWHPNGWTL
YQVLEQYMRKVQRENGYLEIKTPQVVDRSLWEKSGHWANYADNMFTTQSENRDYAIKPMNCPCHVQVF
NQGLKSYRELPMRLAEFGACHRNEPSGALHGIMRVRGFTQDDAHIFCTEEQMQAESAAFIKLTMDVYR
DFGFTEVEMKLSTRPEKRVGSDELWDRAEAALAAALDSAGLAYDLQPGEGAFYGPKIEFSLKDCLGRV
WQCGTLQLDFNLPIRLGAEYVCEDNSRKHPVMLHRAILGSFERFVGILIEHYEGAFPAWLAPTQAVIM
NITDKQADFAAEVEKTLNESGFRAKSDLRNEKIGFKIREHTLLKVPYLLVIGDREVEMQTVAVRTREG
ADLGSMPVAQFAEFLAQAVSRRGRPDSE
